# Force-induced changes of PilY1 drive surface sensing by *Pseudomonas aeruginosa*

**DOI:** 10.1101/2021.08.24.457478

**Authors:** Shanice S. Webster, Marion Mathelié-Guinlet, Andreia F. Verissimo, Daniel Schultz, Albertus Viljoen, Calvin K. Lee, William C. Schmidt, Gerard C. L. Wong, Yves F. Dufrene, George A. O’Toole

**Affiliations:** Department of Microbiology and Immunology, Geisel School of Medicine at Dartmouth, Hanover NH 03755, USA; Louvain Institute of Biomolecular Science and Technology, Université Catholique de Louvain, Croix du Sud, 4-5, bte L7.07.07, B-1348 Louvain-la-Neuve, Belgium; Institute for Biomolecular Targeting (bioMT), Geisel School of Medicine at Dartmouth, Hanover NH 03755, USA; Department of Bioengineering, University of California Los Angeles, CA 90095; Department of Chemistry and Biochemistry, University of California Los Angeles, CA 90095; California NanoSystems Institute, University of California Los Angeles, CA 90095

**Keywords:** type 4 pili, force, PilY1, von Willebrand A domain, surface sensing, c-di-GMP

## Abstract

During biofilm formation, the opportunistic pathogen *Pseudomonas aeruginosa* uses its type IV pili (TFP) to sense a surface, eliciting increased second messenger production and regulating target pathways required to adapt to a surface lifestyle. The mechanisms whereby TFP detect surface contact is still poorly understood, although mechanosensing is often invoked with little data supporting this claim. Using a combination of molecular genetics and single cell analysis, with biophysical, biochemical and genomics techniques we show that force-induced changes mediated by the von Willebrand A (vWA) domain-containing, TFP tip-associated protein PilY1 are required for surface sensing. Atomic force microscopy shows that PilY1 can undergo force-induced, sustained conformational changes akin to those observed for mechanosensitive proteins like titin. We show that mutation of a single cysteine residue in the vWA domain results in modestly lower surface adhesion forces, increased nanospring-like properties, as well as reduced c-di-GMP signaling and biofilm formation. Mutating this cysteine has allowed us to genetically separate TFP function in twitching from surface sensing signaling. The conservation of this Cys residue in all *P. aeruginosa* PA14 strains, and its absence in the ~720 sequenced strains of *P. aeruginosa* PAO1, could contribute to explaining the observed differences in surface colonization strategies observed for PA14 versus PAO1.

**Importance:** Most bacteria live on abiotic and biotic surfaces in surface-attached communities known as biofilms. Surface sensing and increased levels of the second messenger molecule c-di-GMP are crucial to the transition from planktonic to biofilm growth. The mechanism(s) underlying TFP-mediated surface detection that triggers this c-di-GMP signaling cascade are unclear. Here, we provide a key insight into this question: we show that the eukaryotic-like, vWA domain of the TFP tip-associated protein PilY1 responds to mechanical force, which in turn drives production of a key second messenger needed to regulate surface behaviors. Our studies highlight a potential mechanism that could account for differing surface colonization strategies.

## Introduction

*Pseudomonas aeruginosa* is a ubiquitously distributed opportunistic pathogen that encounters mechanical forces during surface sensing – a crucial first step for biofilm formation. The type four pili (TFP) motility appendage is integral to surface sensing and is thought to transduce a force-induced signal to the cell interior by detecting the resistance to retraction when cells are surface engaged [1], activating the production of cAMP and c-di-GMP, and regulating target genes that control biofilm formation [2–4]. While the importance of the TFP and its tip associated protein, PilY1, in surface sensing has been proposed, direct evidence of how the TFP/PilY1 sense the surface is lacking. Indeed, much of the supporting evidence of a role for this appendage as a key surface sensor is deductive, or alternatively, rely on biological responses or phenotypic changes that are observed during the switch from planktonic to surface-attached growth. In this study, we thus take a multi-disciplinary approach to investigate the mechanism whereby the TFP via the tip-associated protein, PilY1, is directly involved in surface sensing.

PilY1 is part of the priming complex together with the minor pilins that facilitate incorporation of the PilA subunits into the base of the growing pilus fiber during elongation [5, 6]. During polymerization, the minor pilins and PilY1 are pushed to the tip of the growing pilus. PilY1 has a C-terminal domain that resembles PilC from *Neisseria gonorrhoea* and a N-terminal von Willebrand A (vWA) domain (Fig. 1A) that is structurally similar to the A2 domain of the human von Willebrand factor (vWF), a force sensing glycoprotein important in stopping bleeding [4]. The vWA domain of PilY1 has the classical Rossman fold – central beta sheets surrounded by amphipathic alpha helices [7] – and a perfectly conserved metal ion dependent adhesion site (MIDAS) containing the conserved DxSxS…T…F motif [8]. vWA domains have been reported in TFP-associated proteins from other organisms. For example, the vWA domain of the major pilin in *Streptococcus agalactiae* is essential for adhesion [9] and the MIDAS motif in the vWA domain of the major pilin in *Streptococcus sanguinis* has recently been shown to be important in binding to eukaryotic cells [10]. Like the vWF [11], the vWA domain of *P. aeruginosa* PA14 PilY1 protein also has a high number of cysteine residues; seven out of the 11 cysteines in PilY1 are in its vWA domain. Interestingly, during vascular damage, when exposed to high shear forces due to blood flow, the vWF transitions from a globular to a stretched conformation [12, 13]. This transition is thought to be mediated by a disulfide bond switch exposing specific sites that allow platelets to bind [14–16]. Thus, vWF cysteine residues, depending on their redox state, are key to force sensing, a property that could be hypothesized for cysteine residues in the vWA of PilY1.

**Figure 1.**
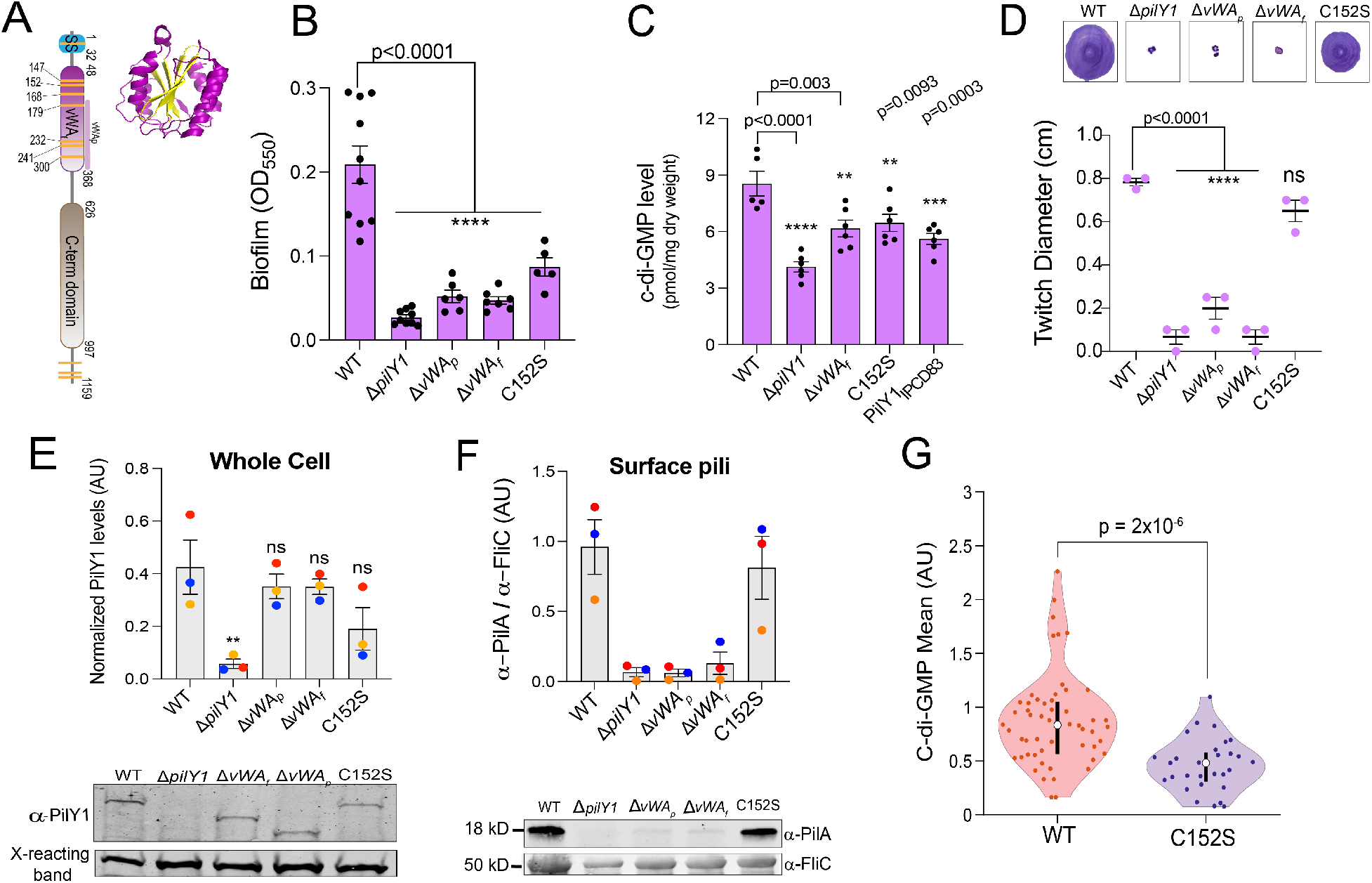
The von Willebrand A (vWA) domain and Cys152 residue of PilY1 are important for regulating c-di-GMP levels and biofilm formation. **A.** Schematic showing domain organization of the PilY1 protein. The signal sequence (SS – blue, amino acids 1-32), vWA domain (pink, amino acids 48-368) and C-terminal domain (brown, amino acids 626-997) are highlighted. vWA_p_ (amino acids S168-S365) denotes a portion of the vWA domain that is deleted in a mutant analyzed in the subsequent panels. Yellow stripes represent the cysteines residues present in the protein. The vWA domain contains seven of the 11 cysteine residues present in the full length PilY1 protein with the SS and the C-terminal region having one and three cysteine residues, respectively. Inset: Ribbon diagram showing the vWF A2 domain (PDB 3GXB). The domain shows a classical Rossmann fold [7], comprised of central β-sheets (yellow) surrounded by a-helices (purple). **B.** Biofilm formation measured at OD_550_ for WT, the Δ*pilY1* deletion mutant, the vWA variants, and the Cys152S mutant in a static 96 well biofilm assay performed in M8 medium salts plus supplements (see Materials and Methods) and incubated at 37 °C for 24 h. vWA_p_ (amino acids 178-365, see panel A) and vWA_f_ indicate a partial and full deletion (amino acids 48-368) of the vWA domain, respectively. Data are from at least five biological replates each with eight technical replicates. **C**. Quantification of global c-di-GMP levels by mass spectrometry for WT and the indicated mutants shown in picomole per milligram dry weight. Cells were grown on 0.5% agar plates prepared with M8 medium salts plus supplements, then scraped from the plates after incubation for 37 °C for 14-16 h. Data are from six biological replicates each with two technical replicates. **D**. Twitch diameter (cm) for WT and the indicated mutants measured after inoculating LB plates from overnight cultures, then incubating the plates for 24 h at 37 °C plus an additional day at room temperature. Representative images of twitch zones are shown above the graph. Data are from three biological replicates. **E**. Quantification of normalized PilY1 protein levels in whole cell (in arbitrary units (AU)) for WT and the indicated mutants. Cells were sub-cultured from an overnight culture and grown to mid-log phase in M8 medium salts plus supplements and normalized to the same OD_60_0 value. Protein levels in whole cell extracts are normalized to a cross-reacting band at ~60 kDa, which is used as an additional loading control. The Cys152S mutant shows a modest but not significant reduction in level in whole cell extracts. A representative Western blot image for PilY1 and the cross-reacting band are shown below the graph. **F.** Quantification of normalized surface pili levels. PilA (~18 kDa) protein levels are used as a surrogate for surface pili levels, which are normalized to levels of the flagellar protein, FliC (~50 kDa). A representative Western blot is shown below the graph. All Western blot data are from three biological replicates in three independent experiments. Dots with the same color represent the same biological replicate; different colors indicate different biological replicates. p values: p ≤ * 0.05, ns, not significant. All error bars in Figure 1 are standard error of the mean (SEM) and statistical significance was determined by one-way ANOVA and a Dunnett’s post-hoc test, p-values: p ≤ **** 0.0001, p ≤ *** 0.001, p ≤ ** 0.01, ns, not significant. **G.** Violin plots showing the mean c-di-GMP of the WT strain and a strain expressing the vWA-Cys152S PilY1 variant during early biofilm formation. c-di-GMP level was quantified from GFP intensity determined on a cell-by-cell basis in a microfluidic chamber carrying the P*_cdrA_*-GFP construct, which is a reporter of c-di-GMP levels. NOTE: The WT data shown here was first reported in a previous publication [20]; each strain analyses is done independently, in the same system and medium, with the same microscope at identical settings and processed as reported [20]. Given that each analysis is independent but performed identically, we can compare data from previous studies. Each data point represents one tracked cell through an entire division cycle. Statistical significance was determined using the Kruskal-Wallis test, p = 2×10^−6^.

Although PilY1 is known to be important in responding to shear forces and in increasing c-di-GMP levels [3, 17, 18], the precise role of its vWA domain in surface sensing and c-di-GMP signaling is unclear. Our previous genetic studies show that PilY1 and the vWA domain are important for surface-dependent stimulation of c-di-GMP production [3, 18]. These studies also showed that while the C-terminal domain of PilY1 was dispensable for surface-dependent c-di-GMP production, strains with mutations in the vWA domain failed to regulate c-di-GMP levels and c-di-GMP-related behaviors [18]. Additionally, deletion of the vWA domain is shown to lock PilY1 in a constitutively active signaling conformation that induces virulence independent of surface attachment [4], suggesting multiple roles for the vWA domain in the surface-attached biology of *P. aeruginosa*.

Recent cryo-electron tomography studies show the vWA domain of PilY1 to be situated at the very tip of the pilus fiber [5] indicating that this domain is likely in immediate contact with the surface and therefore could be directly engaged in surface sensing. Based on the similarities between the human vWF and the vWA domain of PilY1, and its importance in downstream c-di-GMP signaling, we hypothesized that force-induced conformational changes originating from the vWA domain of PilY1 are mediated by conserved cysteine residues within this putative mechanosensing domain, and together these features of PilY1 are critical for surface sensing. We explore these hypotheses here.

## Results

### The von Willebrand A (vWA) domain of PilY1 regulates c-di-GMP levels and biofilm formation

To address the role of the vWA domain of PilY1 in surface sensing and c-di-GMP signaling, we made chromosomal deletions that removed a part (Δ*vWA_p_*) or the full (Δ*vWA_f_*, **Fig. 1A**) vWA domain, then performed static biofilm assays and measured global levels of c-di-GMP (**Fig. 1B** and **C**). Our bulk assays show that both the Δ*vWA_p_* and Δ*vWA_f_* variants resulted in a significant decrease in biofilm formation and reduction in global c-di-GMP levels (as seen for the Δ*vWA_f_* variant) as compared to WT. These vWA variants also resulted in no twitching motility (**Fig. 1D**). To confirm these twitch phenotypes, another more sensitive assay was used based on the lysis of the host cells *P. aeruginosa* PA14 by the lytic DMS3vir phage, which specifically targets the TFP and also requires retraction of surface-expressed pili for infection [19]. Strains carrying the Δ*vWA_p_* and Δ*vWA_f_* variants showed partial zones of clearing in a phage plaquing assays (**Fig. S1A**), indicating that these strains retained some TFP function. To further ensure that the decrease in biofilm formation and reduced c-di-GMP levels were not due to protein instability, we examined steady state levels of the vWA variants in whole cell extracts. Both the Δ*vWA_p_* and Δ*vWA_f_* PilY1 variants were stable and showed a non-significant reduction in whole cell levels as compared to WT (**Fig. 1E**). However, little surface pili could be detected in the strains expressing the Δ*vWA_p_* and Δ*vWA_f_* variants (**Fig. 1F**), which likely explains the lack of full pilus function observed in the twitch assays (**Fig. 1D**). The presence of plaques (**Fig. S1A**), however, indicates that there are some surface pili, a finding consistent with our Western blots (**Fig. 1F**).

**Figure S1.**
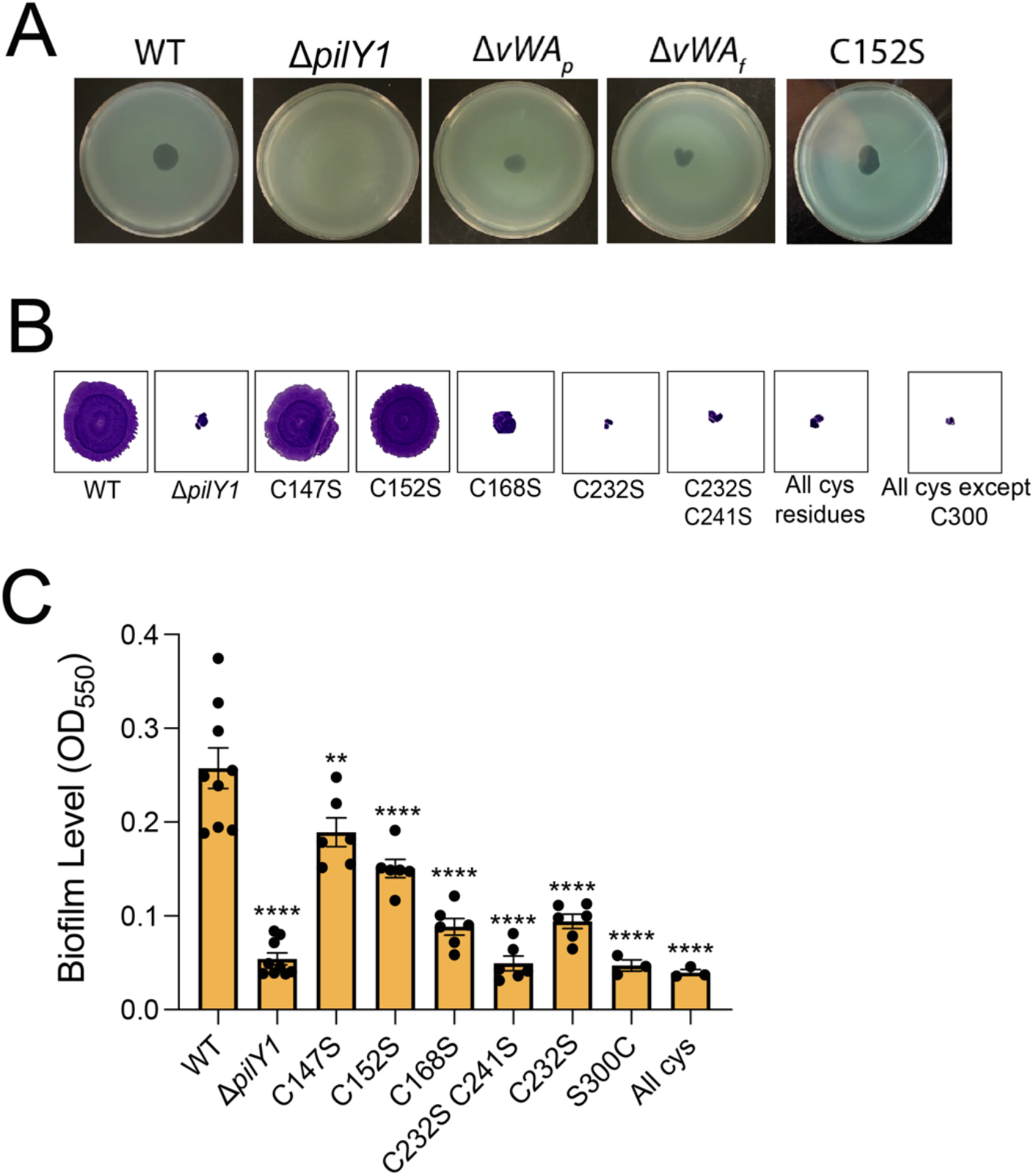
Partial functionality of vWA variants and phenotypic analysis of other cysteine vWA mutants. **A.** Plaquing assay with phage DMS3_*vir*_ versus the WT and the indicated mutants as hosts. Zones of clearing shown for WT and the strain expressing the vWA-Cys152S mutant protein are similar, which indicates a similar degree of TFP function. The Δ*pilY1* mutant serves as the negative control. **B.** Representative images of twitch zones stained with crystal violet shown for WT, the Δ*pilY1* or strains expressing PilY1 variants with point mutations in the Cys residues in the vWA domain following incubation at 37 °C for 24 h plus one additional day at room temperature. Twitching serves as a measure of TFP function. **C.** Biofilm level measured at OD550 for WT and the mutants shown in panel B using the 96 well static biofilm assay after 24 hrs at 37 _C_, as described in the Materials and Methods.

### The Cys152 residue of the vWA domain is important for promoting biofilm formation and regulating c-di-GMP levels

Multimerization and conformational changes required for function in blood clotting by the human vWF are mediated by cysteine residues [15, 16, 21]. Shear forces due to blood flow during vascular damage have been shown to induce disulfide bond cleavage, which results in the protein adopting a new, stretched conformation [21, 22]. Inspired by these studies and the high number of cysteines in the vWA domain of PilY1 (**Fig. 1A**), we hypothesized that one or more cysteines in the vWA domain of PilY1 might be important for mediating conformational changes in PilY1 and/or the pilus fiber that could in turn impact surface sensing and downstream c-di-GMP signaling. To test this hypothesis, we performed targeted mutagenesis of the cysteine residues in the vWA domain of PilY1 with the aim of identifying one or more of these residues that impact biofilm formation but still retain TFP function as assessed by twitching assays. In all cases, the mutations were introduced into the chromosomal copy of the *pilY1* gene, thus the mutants were expressed under the native *pilY1* promoter and in their native chromosomal context. Of the seven individual and combination cysteine residues mutated, five resulted in decreased biofilm formation but no twitching motility (**Fig. S1B and S1C**). However, two residues, when mutated (vWA-C147S and vWA-Cys152S) displayed decreased biofilm formation but retained twitching motility (**Fig. 1B, D** and **Fig. S1B**). Because the vWA-Cys152S mutation yielded the stronger biofilm phenotype, we focused on this mutant for all subsequent analyses.

We next measured c-di-GMP levels globally and on a cell-by-cell basis for the strain expressing the vWA-Cys152S variant. Compared to WT, the vWA-Cys152S mutant showed significantly reduced levels of c-di-GMP based on bulk measurements of cell extracts and on a single-cell basis (**Fig. 1C** and **Fig. 1G**, respectively). Note: the WT data shown in the single cell data (**Fig. 1G**) was first reported in a previous publication [20]. Analyses of WT and vWA-Cys152S were done independently, in the same system and medium, analyzed with the same microscope at identical settings and processed as reported [20]. Given that each investigation is independent but performed identically, it allows us to compare data from this previous report.

Given the similar levels of twitching motility for the strain carrying the vWA-Cys152S mutant and the WT, we predicted that this point mutation would yield a stable PilY1 protein. Western blot studies of whole cells showed that the vWA-Cys152S variant is stable and shows a modest but non-significant reduction in protein levels as compared to WT PilY1 (**Fig. 1E**). Additionally, surface pili levels for the strain expressing the vWA-Cys152S mutant protein are comparable to WT (**Fig. 1F**). These results are consistent with the vWA-Cys152S mutant showing similar levels of twitching motility (**Fig. 1D**) and plaque formation (**Fig. S1A**) compared to WT. Of note, none of the observed phenotypes are due to differences in growth rates as the vWA-Cys152S strain along with all vWA mutants used in this study have the same growth kinetics as WT (**Fig. S2**).

**Figure S2.**
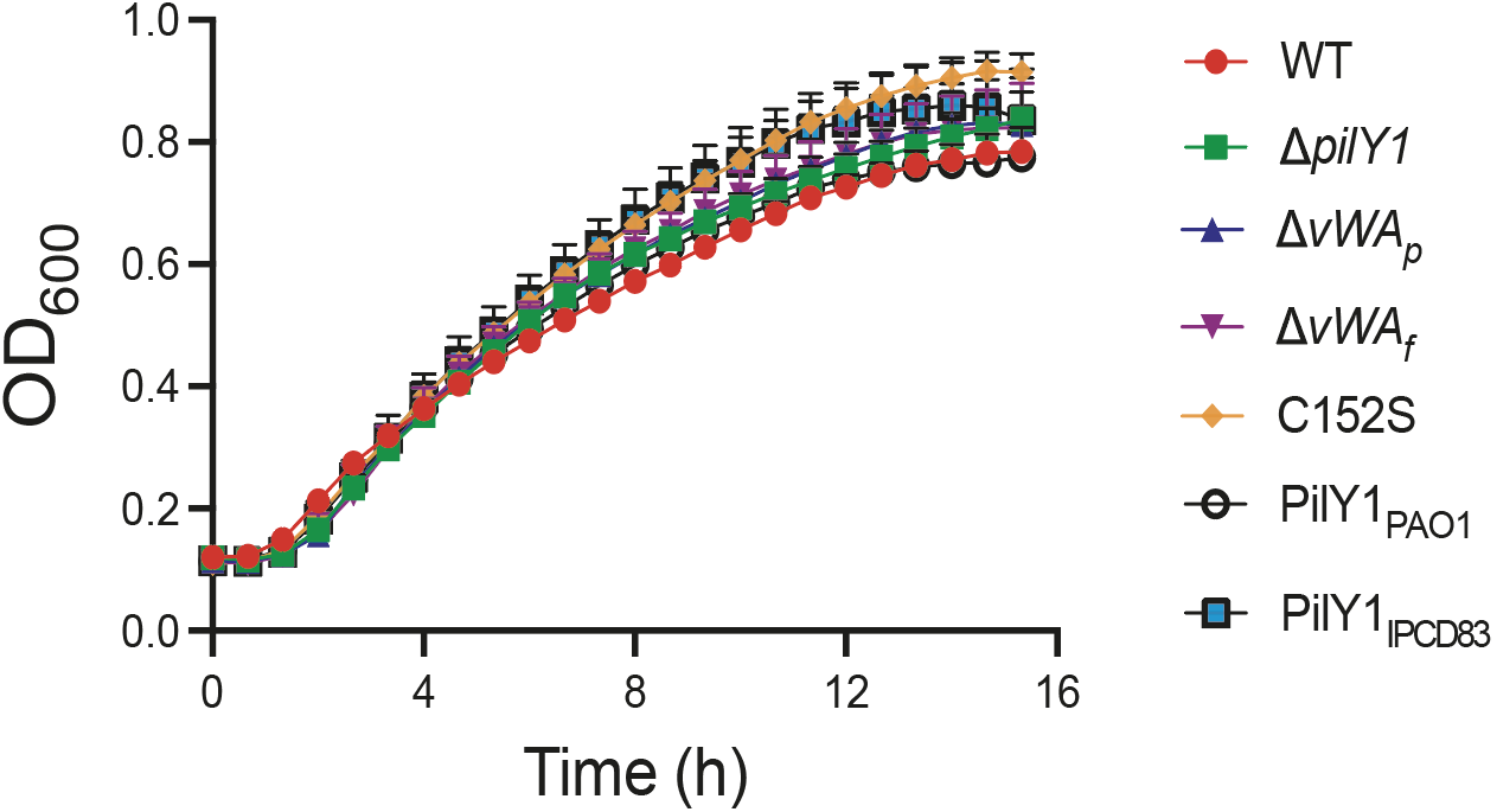
Growth curves for WT and the strains expressing the PilY1 variants. Growth assays were performed in M8 minimal salts medium supplemented with casamino acids, glucose and magnesium sulfate. This medium was also used for all macroscopic biofilm assays, c-di-GMP measurements, plaquing assays and AFM studies. The data are from three biological replicates each with two technical replicates. There is no significant difference among the growth kinetics of each strain. Error bars show SEM and statistical significance was determined at each time point using one-way analysis of variance (ANOVA) using multiple comparisons test.

### The vWA-Cys152S variant of PilY1 is associated with lower surface adhesion forces and altered force-induced behaviors

In light of the key role of the vWA domain in biofilm formation and c-di-GMP regulation, we next sought to investigate the different surface adhesion behaviors of *P. aeruginosa* strains expressing WT PilY1, or the PilY1 variants with the Δ*vWA_f_* or the vWA-Cys152S mutations. To this end, we used atomic force microscopy (AFM), a powerful multifunctional technique that has been instrumental in deciphering the adhesion and nanomechanical properties of bacterial pili, at the single-cell and single-molecule levels [23–25]. More specifically, we recorded the force experienced by a hydrophobic AFM tip when probed against the TFP of surface engaged bacterial cells as a function of the tip-sample surface distance (**Fig. 2A**). From the resulting force-distance curves, binding probability and adhesion forces were determined on multiple living cells. As illustrated in the representative force histograms (**Fig. 2B**), the vWA-Cys152S mutant showed a lower adhesion force than WT cells (F = 133 ±89 pN and 211 ±72 pN respectively, p<0.001), indicating that the Cys152S mutation impacts the interaction strength. However, both WT and vWA-Cys152S PilY1 cells showed a similar binding probability to the hydrophobic AFM tip (**Fig. 2C**), a result that is consistent with both strains having similar levels of surface pili (**Fig. 1G**). Cells with the full deletion of the vWA domain (ΔvWA_f_) showed an ~0% binding probability (**Fig. 2C**) to the hydrophobic tip and a low adhesion force (~45 pN, **Fig. 2B**), likely due to a low number of surface pili (**Fig. 1F**). These data suggest that the observed force curves are dependent on the TFP-associated PilY1.

**Figure 2.**
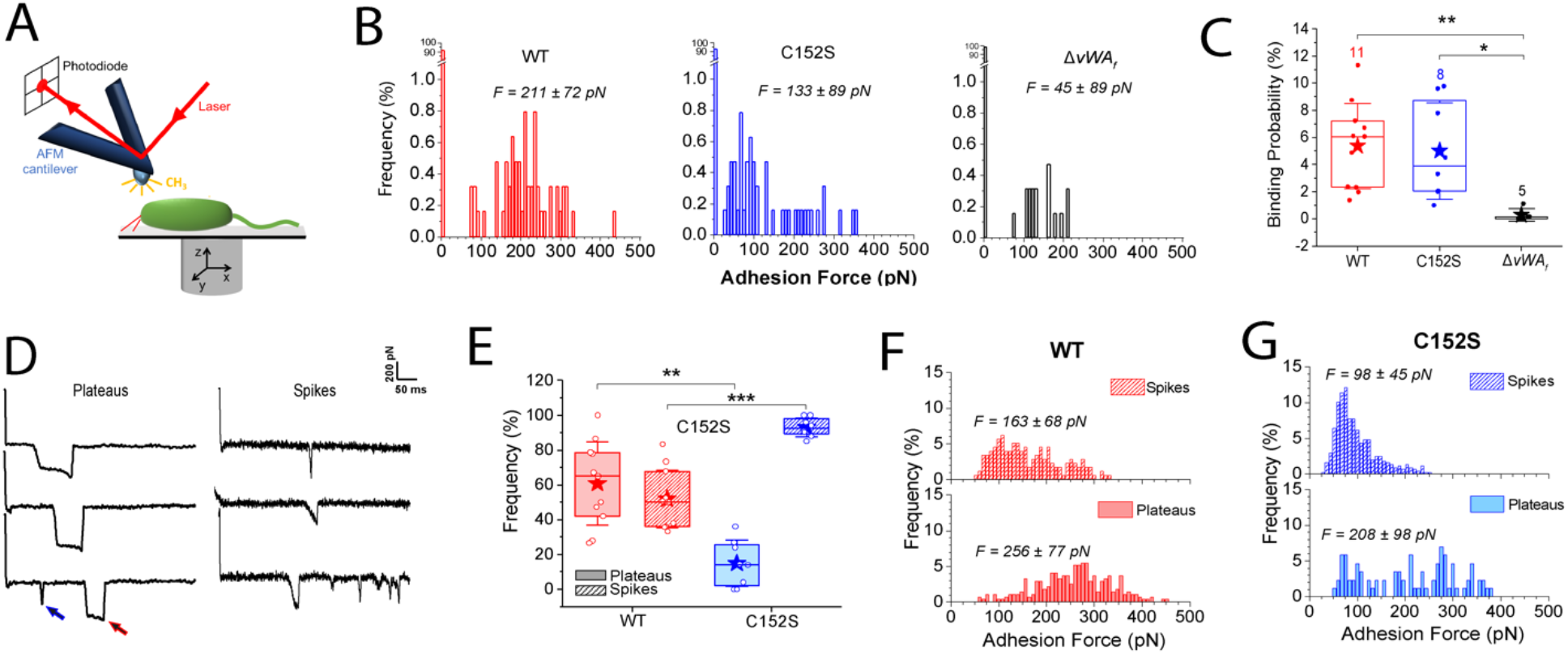
Strains expressing the PilY1-Cys152S mutation display less adhesion force and altered mechanical behaviors compared to strains expressing the WT PilY1. **A.** Scheme of the AFM setup showing that piliated *P. aeruginosa* is probed with a hydrophobic AFM tip at the free end of the AFM cantilever. Adhesive interactions occurring between the pilus/cell body and the AFM tip cause a deflection of the cantilever, directly proportional to force, which is recorded by a laser beam focused at the AFM tip’s free end and reflected back to a photodiode. **B.** Adhesion force histograms between the hydrophobic AFM tip and a representative WT strain, or strains expressing the Cys152S or Δ*vWA_f_* variants of PilY1. For WT: 211 ± 72 pN (n = 55 adhesive curves); for the vWA-Cys152S: 133 ± 89 pN (n = 47) and for the ΔvWA_f_: 45 ± 89 pN (n = 16). **C.** Box plots comparing the binding probability of cells expressing the WT PilY1 or of strains expressing the Cys152S or Δ*vWA_f_* variants of PilY1 are shown. The number of probed cells is indicated. Stars are the mean values, lines the medians, boxes the 25-75 % quartiles and whiskers the standard deviation (SD). Student t-test: * p ≤ 0.05, ** p ≤ 0.01. **D.** Representative retraction force profiles exhibited by the WT or Cys152S mutant cells sorted based on their shape. Plateaus are defined as adhesive events with a “step” behaviour, i.e., a constant sustained force over a defined length of time while spikes are defined as sharp adhesive events with a single minimum. A single retraction profile can feature several plateaus (red arrow), spikes (blue arrow) and even both signatures can occur as marked by the arrows. **E.** Box plots comparing the occurrence of plateaus (shaded) or spikes (striped) signatures for the WT and Cys152S mutant cells. The number of probed cells is as described in panel C. For the WT, plateaus = 60.8 +/− 24.0 % and spikes = 51.9 +/− 16.6 %, *n =* 11, and for Cys152S, plateaus = 14.9 +/− 13.3 % and spikes = 93.1 +/− 5.4 %, *n =* 8. Stars are the mean values, lines the medians, boxes the 25-75% quartiles and whiskers the SD. Student t-test: ** p ≤ 0.01, *** p ≤ 0.001. **F** and **G.** Distribution of the adhesion forces exhibited by either the plateaus or the spikes for the WT **(F)** or the strain carrying the Cys152S mutant of PilY1 (**G**). The mean values are provided along with the histograms. All data were obtained by recording force-distance curves in medium containing M8 salts with an applied force of 250 pN and a pulling speed of 5 *μ*m/s at room temperature.

For cells expressing the WT PilY1 and the vWA-Cys152S variant, which both showed adhesion to the hydrophobic AFM tip, two distinct adhesive behaviors were observed, plateaus (red arrow) and spikes (blue arrow; **Fig. 2D**). Plateaus are defined as adhesive events with a “step” behavior, that is, a constant sustained force over a defined length of time, while spikes are defined as sharp adhesive events with a single minimum and are reflective of a nanospring behavior [25]. Plateaus and spikes are not mutually exclusive in their appearance and frequency. Cells expressing WT PilY1 or the vWA-Cys152S variant showed plateaus and spikes, however, the frequency of these behaviors differed significantly between the strains (**Fig. 2E**). Cells expressing the WT PilY1 had a similar proportion of plateaus (~61%) and spikes (~52%; the sum can be >100% because some force curves can have both features). In contrast, cells expressing the vWA-Cys152S mutant of PilY1 showed a significantly lower frequency of plateaus (~15% compared to ~61% for the WT) and a much higher frequency of spikes (~93% compared to ~52% for the WT; **Fig. 2E**). These data indicate that a single point mutation in the PilY1 vWA domain can have a marked impact on the cell’s mechanical behavior.

Finally, the magnitude of the adhesive signatures for both spikes and plateaus were higher for cells with WT PilY1 than those cells expressing the vWA-Cys152S variant (**Fig. 2F** and **G**), consistent with the observation that cells expressing WT PilY1 can sustain globally higher adhesive forces than the cells expressing the vWA-Cys152S mutant (**Fig. 2B**). Interestingly, for both strains the observed plateau forces are higher than those observed for the spikes, which, along with the higher frequency of plateaus observed in WT PilY1, also explains the higher forces sustained by the WT cells.

Our data above indicate that the observed adhesive forces as well as the plateau and spike signatures observed for strains expressing WT PilY1 protein versus the vWA-Cys152S mutant protein were dependent PilY1 and its vWA domain. We next asked where these force profiles were dependent of the TFP. Because PilY1 is cell-surface-associated and can be secreted to the cell-surface independent of the TFP machinery [3], we expressed plasmid-borne WT PilY1 and the vWA-Cys152S mutant protein in a Δ*pilA* background, lacking the full pilus fiber, and performed AFM experiments. Both strains expressing the WT PilY1 protein and the vWA-Cys152S mutant protein showed little adhesion to the hydrophobic tip (binding probability < 1%; **Fig. S3A-C**). The scarce adhesive events recorded for the strains expressing these proteins in the Δ*pilA* background were significantly lower than those exhibited when the pilus was present, and plateau signatures were never observed (**Fig. S3D**). Instead, typical receptor-ligand signatures were recorded, resembling a spike signature, but with very short rupture length consistent with the length of the protein that is stretched while the AFM tip retracts away from the bacterium (**Fig. S3D**). Together, the genetic and AFM data support the hypothesis that the adhesive forces measured, as well as the plateaus and spikes signatures exhibited by the strains expressing the WT PilY1 protein and the vWA-Cys152S mutant protein, are due to both PilY1 plus the pilus fiber.

**Figure S3.**
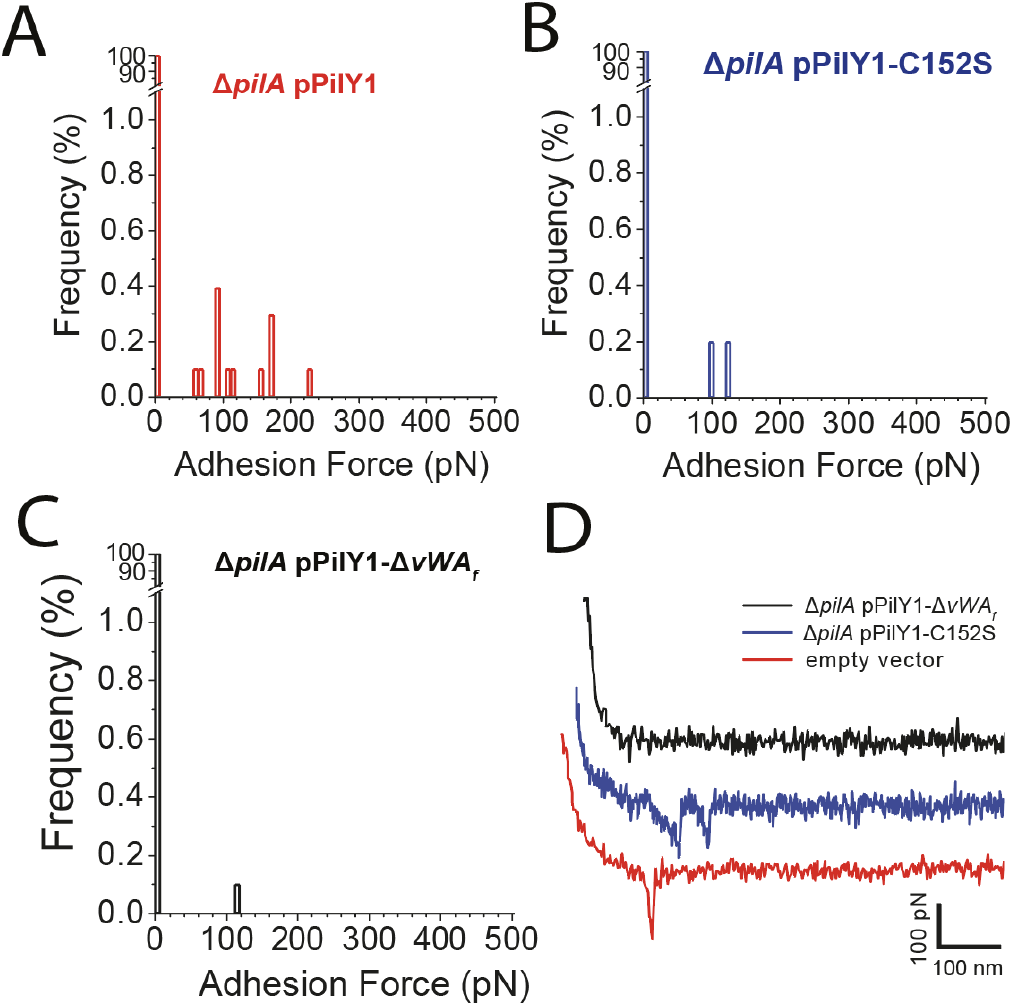
The pilus fiber is required for adhesion to a surface. **A-C.** Adhesion force histograms obtained by recording force-distance curves between the hydrophobic cantilever tip and representative Δ*pilA*/pPilY1 (**A**), Δ*pilA*/pPilY1-Cys152S (**B**) and Δ*pilA*/pPilY1-ΔvWAf (**C**) cell. **D.** Representative retraction force profiles shown for the same strains.

### The vWA-Cys152S mutation has a negligible impact on the solution conformation of the vWA domain

Given our findings of the significant difference in mechanical behaviors observed for the strains expressing the WT PilY1 protein and the vWA-Cys152S mutant protein when these strains are engaged with a surface and thus under mechanical tension, we next determined whether this single cysteine mutation affected the conformation of purified, isolated vWA domain of PilY1 in solution. We focused on the vWA domain because despite attempts with several different expression systems, we were unable to purify stable, full-length PilY1 or the N-terminal domain of this protein. We cloned WT vWA and vWA-C152S domains (aa 30-369) as glutathione-S-transferase (GST) fusion proteins to enhance stability and facilitate purification. A GST domain and a HRV-3C protease cleavage site were added to the N-terminus of vWA and the resulting fusion proteins were overexpressed in *E. coli* cells and purified to homogeneity (**Fig. 3A**). The HRV-3C protease cleavage site was confirmed by sequencing. Unfortunately, repeated attempts to efficiently cleave the GST domain from the vWA proteins with protease HRV 3C were unsuccessful, perhaps due to steric occlusion of the protease binding site in the purified proteins. Thus, the studies below were done using GST-vWA fusion proteins.

**Figure 3.**
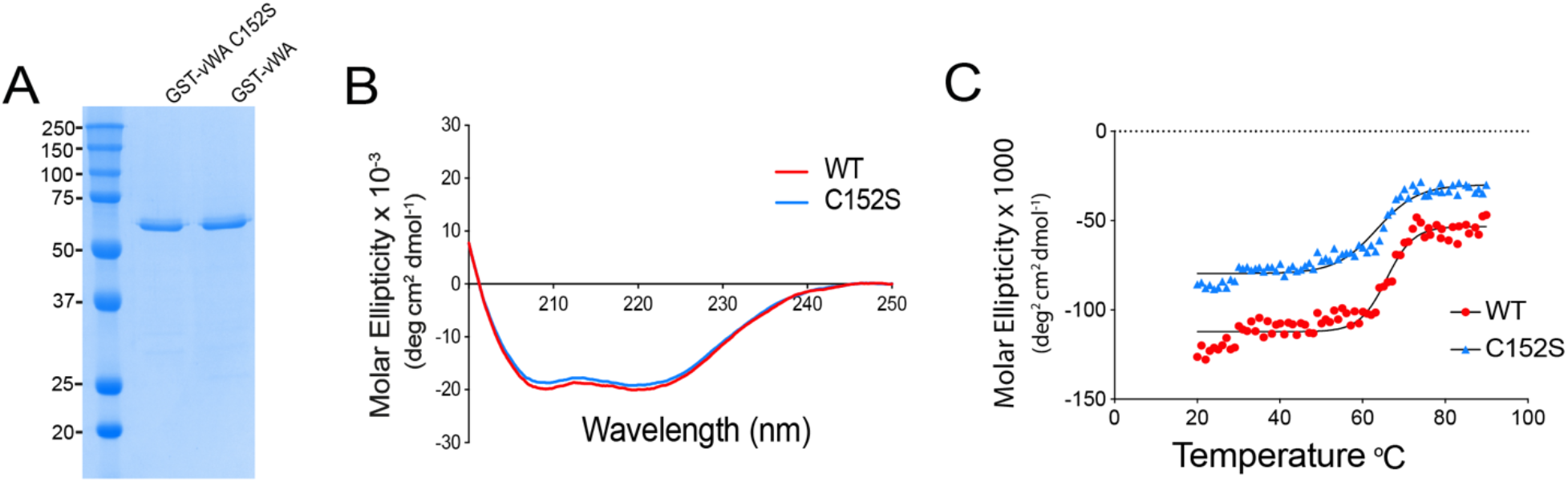
vWA-C152S mutation does not substantially alter conformation of the vWA domain. **A.** Coomassie Blue-stained SDS-PAGE of ~ 1 μg of purified wild-type GST-vWA and GST-vWA-C152S fusion proteins expressed from a pGEX plasmid backbone and purified from *E. coli* BL21-DE3 cells as detailed in the Materials and Methods. The molecular weight markers are indicated. **B.** Far-UV Circular dichroism (CD) spectra shown in molar ellipticity for the WT GST-vWA (red line) and GST-vWA-C152S mutant (blue line) between 195 and 250 nm at 20 ° C. **C**. Curves of ellipticity at 208 nm wavelength as a function of temperature for WT and mutant fusion proteins. Spectra were recorded for each sample from 20 to 90 ° C in 1 ° increments. Curves were fitted to a Boltzmann sigmoidal equation and the V_50_ value was determined (65.8 versus 63.5 °C for GST-WT-vWA and GST-vWA-C152S fusion variant, respectively).

We performed far-UV circular dichroism (CD) spectroscopy to determine the secondary structure of the WT and mutant fusion proteins and to assess the thermal stability of WT-vWA and the vWA-Cys152S variants (**Fig. 3B** and **3C**). Far-UV CD spectra of the GST-WT-vWA and GST-vWA-Cys152S fusion proteins were monitored at wavelength scans between 195 and 250 nm. Both WT and mutant spectra showed the presence of two distinct negative peaks centered at 208 and 222 nm, typical of α-helical proteins (**Fig. 3B**). Overall, the dichroic spectra for GST-WT-vWA and GST-vWA-Cys152S were similar. Measuring CD as a function of temperature can be used to determine the effects of mutations on protein stability. Analysis of the ellipticity curves in the range of 20 to 90 °C showed the melting temperatures of GST-WT-vWA and GST-vWA-C152S fusion variant to be similar (65.8 versus 63.5 °C; **Fig. 3C**), suggesting that the C152S mutation did not perturb the secondary structure of the domain in solution (i.e., in the absence of mechanical force).

### Genomic analyses reveal that PAO1 strains lack the vWA-C147 and vWA-Cys152 cysteine residues that are present in PA14 strains, with associated functional consequences

Given our findings that cells expressing the vWA-Cys152S mutation impact surface sensing, c-di-GMP levels and biofilm formation (**Fig. 1B-C, Fig. S1B**) in *P. aeruginosa* PA14, we analyzed whether the Cys152 residue was conserved across *P. aeruginosa* strains. We leveraged PilY1 sequences from the international *P. aeruginosa* consortium database (IPCD), a repository for thousands of *P. aeruginosa* isolates from a diverse range of environments [26]. We analyzed the phylogenetic relationship of PilY1 amino acid sequences from a total of 852 *P. aeruginosa* genomes and found two distinct clades, PA14 (red dot) and PAO1 (dashed circle; **Fig. 4A**), largely consistent with a previous report by Levesque and colleagues [26]. The PilY1 sequence from the strain, IPCD83 (blue dot), falls within the PAO1 clade. Alignment of the amino acid sequences of the vWA domain of PilY1 from the PA14, PAO1 and the IPCD83 strains show that five of the seven cysteines (magenta) in the vWA domain of PA14 are highly conserved in PAO1 and IPCD83, although the spacing of the residues varies in some cases (**Fig. 4B**). All three domains consist of positive, negative, polar and hydrophobic amino acids shown in red, blue, green and grey, respectively. Of note is the high abundance of polar residues in the vWA domains of all three strains.

**Figure 4.**
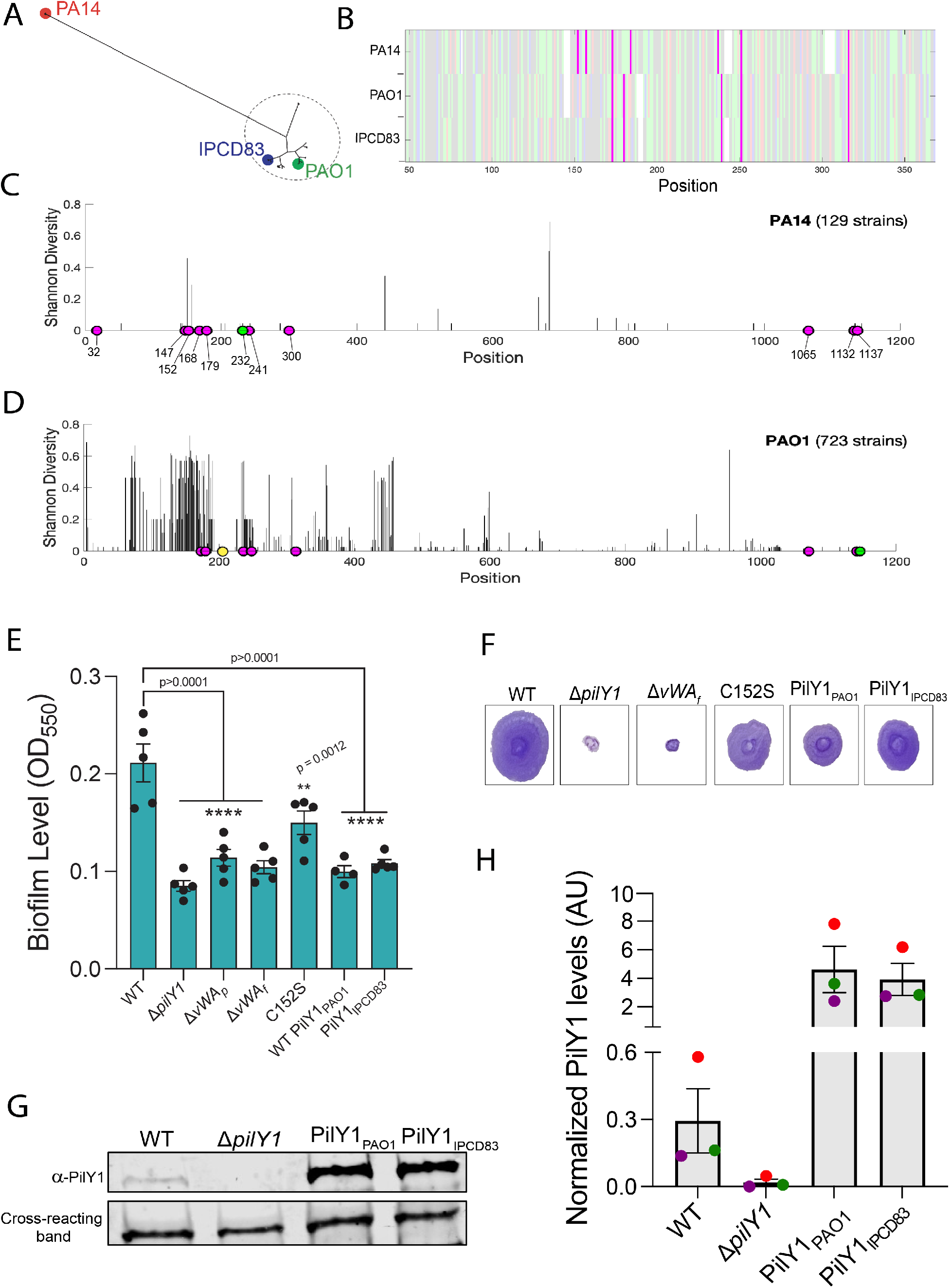
Comparative genomic analyses reveal sequence and functional differences between PA14 and PAO1 alleles of PilY1. **A.** Phylogenetic tree of PilY1 amino acid sequences obtained from the IPCD database of *P. aeruginosa* genomes [26] showing two distinct clades of PilY1 sequences corresponding to strains from the previously determined *P. aeruginosa* PA14 and PAO1 clades. The strain labeled IPCD83 is an isolate within the PAO1 clade. **B**. Alignment of the vWA domain (48 to 368) of PilY1 proteins found in PA14, PAO1 and IPCD83 strains, with cysteines highlighted in magenta. Positive, negative, polar and hydrophobic amino acids are depicted in red, blue, green and grey, respectively. **C**. Shannon diversity index along the PilY1 amino acid sequence for the 129 PilY1 proteins belonging to the PA14 clade. Fully conserved cysteines are highlighted in magenta. One strain is missing the cysteine depicted in green. **D**. Shannon diversity index along the PilY1 amino acid sequence for the 723 versions of PilY1 proteins belonging to the PAO1 clade. Fully conserved cysteines are highlighted in magenta. One strain is missing the cysteine depicted in green, and one strain has an extra cysteine depicted in yellow. **E**. Biofilm formation measured at OD_550_ in a static 96 well assay for the indicated strains. Hybrid *P. aeruginosa* PA14 strains carry the PilY1 protein from PAO1 (PilY1_PAO1_) or the PilY1 protein from IPCD83 strain (PilY1_IPCD83_) replacing the coding region for the *P. aeruginosa* PA14 PilY1 protein. In all cases, the mutant PilY1 variants are expressed from the native locus of *P. aeruginosa* PA14. Error bars are SEM and statistical significance shown was determined by one-way ANOVA and a Dunnett’s post-hoc test. p values: p ≤ **** 0.0001, p ≤ *** 0.001, p ≤ ** 0.01, ns, not significant. **F**. Representative images of twitch zones shown for the indicated strains. **G**. Representative Western blot image for steady state PilY1 protein levels in whole cells WT PilY1, Δ*pilY1*, PilY1 from PAO1 (PilY1_PAO1_) and PilY1 variant from strain IPCD83 (PilY1_IPCD83_). **H.** Quantification of normalized PilY1 protein levels from whole cells for strains shown in G. Protein level is normalized to a cross-reacting band at ~60 kDa. Data are from three biological replicates in three independent experiments. Dots with the same color represent the same biological replicate; different colors indicate different biological replicates.

To examine the amino acid diversity of the PilY1 sequences in the PA14 and PAO1 clades, we computed Shannon diversity index as a measure of sequence diversity (**Fig. 4C** and 4**D**). We aligned PilY1 sequences within the PA14 (**Fig. 4C**) and PAO1 (**Fig. 4D**) clades and calculated Shannon diversity at each amino acid position.

As shown in Figure 4C, there is very little amino acid sequence diversity over the entire PilY1 sequence among the 129 isolates with PA14 versions of PilY1. Interestingly, all strains in the PA14 clade except one contain the 11 cysteines (magenta circles) found in the PA14 reference strain (**Fig. 4C**). Furthermore, each isolate had all seven cysteines in the vWA domain while there was one strain missing vWA-C232 residue (green circle; a residue we found crititcal for TFP-mediated twitching motility, **Fig. S1B**). In contrast to the PA14 clade, strains within the PAO1 clade showed low diversity at the C-terminal domain (amino acid 626-997) and high amino acid diversity in the vWA domain (amino acid 48-368; **Fig. 4D**). Additionally, of the 723 variants of the PilY1 sequences from the PAO1 clade analyzed, only eight cysteines were highly conserved compared to the 11 highly conserved cysteines for the PA14 strains. The vWA domain of the PAO1 clade contains five of the seven conserved cysteines found in the PA14 clade. Interestingly, vWA-147 and vWA-Cys152 residues are not present in any of the PAO1 strains, including the IPCD83 isolate. Recall, that we showed that both vWA-C147 and vWA-Cys152 residues are important in c-di-GMP signaling, and mutations in these residues resulted in strains with decreased biofilm formation but retaining twitching motility (**Fig. 1B** and **Fig. S1B**).

Given the biofilm phenotype of the strain expressing the vWA-Cys152 variant of PilY1 (**Fig. 1B** and **Fig. S1B**) and the role of PilY1 in early biofilm formation and c-di-GMP signaling, we expected that loss of the vWA-Cys152 residue in strains from the PAO1 clade, including IPCD83, should result in similar phenotypes. To test this hypothesis, we cloned the *pilY1* gene from the IPCD83 isolate (PilY1_IPCD83_) or the WT PAO1 strain (WT PilY1_PAO1_) into the native locus of the reference PA14 strain and performed static biofilm assays. Like the vWA-Cys152S mutation, both PAO1 variants expressed in the PA14 strain resulted in significantly decreased levels of biofilm formation as compared to WT (**Fig. 4E**). Quantification of c-di-GMP levels for PilY1_IPCD83_ showed a significant decrease in c-di-GMP level (**Fig. 1C**). Additionally, both the PilY1_PAO1_ and PilY1_IPCD83_ variants still supported twitching motility at a level that is similar to the vWA-Cys152S mutant protein (**Fig. 4F**). The PilY1_PAO1_ and PilY1_IPCD83_ variants showed levels of PilY1 expression that exceed the WT (**Fig. 4G** and **Fig. 4H**), indicating that the observed phenotypes were not due low-level expression of these variants.

## Discussion

Our data show that force-induced changes mediated by one or more cysteine residues in the vWA domain of the TFP tip-associated protein, PilY1, are required for surface sensing and downstream c-di-GMP signaling and biofilm formation. The concept of mechanical force inducing protein conformational changes, that these changes are modulated by disulfide bonds and that such changes in conformation are required for function is well studied in the eukaryotic proteins, titin and vWF. Titin undergoes cycles of folding and refolding that allows it to function as a molecular spring during cycles of muscle relaxation and contraction, respectively [27, 28]. When force is applied, the immunoglobin (Ig) domains of titin unfold and extend [29]. Similarly, increased shear forces due to blood flow cause the vWF to transition from a globular to a stretched conformation [30]; this stretched conformation allows the vWF to bind to platelets and form a clot at sites of vascular damage [31]. Furthermore, the folding and refolding events observed for titin and vWF are mediated by disulfide bonds [32, 33]. For titin, oxidation of the disulfide bond greatly increases both its speed and magnitude of folding [34] while the redox state of the disulfide bond in the A2 domain of the vWF determines exposure of platelet binding sites [21]. Additionally, disulfides bonds in FimH, the adhesin on the type-I pilus in *E. coli* [35], are essential for adhesion under high flow environments [36].

The vWA domain of PilY1 in *P. aeruginosa* PA14 has seven cysteine residues. Our genetic analyses show that two of these residues, vWA-Cys152 and to a lesser extent vWA-Cys147, are critical for PilY1-dependent surface signaling and biofilm formation. Our AFM studies support the conclusion that strains expressing the vWA-Cys152S mutant results in cells that are still capable of surface attachment at the same frequency as the WT, and furthermore, this mutation does not destabilize the PilY1 protein. Using AFM, we show that the WT cells display spike signatures, which are typical of nanospring behaviors [25]. That is, T4P/PilY1 can display elastic properties upon the application of force, but once the force is removed, the pilus rapidly returns to its original conformation. Based on previous work [25] and our data here, these force profiles appear to require both TFP and PilY1. Such nanospring properties are also observed for SpaC, a vWA domain-containing protein that is a key pilus-associated adhesin of *Lactobacillus rhamnosis GG* [23]. Under high mechanical forces, SpaC is shown to behave like a spring. This spring-like behavior is thought to allow the bacterium to withstand higher forces under shear stress when the pilus is stretched, and presumably allow the pilus to engage the surface under strain without snapping [23].

The WT *P. aeruginosa* PA14 strain also shows plateau signatures. One interpretation of these plateau signatures is that they reflect the pilus being bound to the surface at multiple points followed by successive desorption of the pilus [37]. Alternatively, plateaus signatures may be indicative of sustained protein conformational changes. In either case the plateaus observed for WT cells produce high adhesive force signatures, thus likely helping to promote surface engagement.

We found that mutating the Cys152 residue of the vWA domain of PilY results in a reduction in biofilm formation and lower levels of c-di-GMP production. A strain expressing this mutant variant also shows significant changes in mechanical properties (detailed below) when the cell is subjected to force. That these changes in mechanical behavior are dependent on applying a force is in line with our CD and melting curve data which, show no differences in the overall global and secondary structures for WT and the Cys152Ser variant when in solution (i.e., in the absence of an applied force).

The findings from our AFM analysis of the WT and vWA-Cys152Ser allele of PilY1 raise some interesting implications. The ~50-50% distribution of plateaus and spikes observed in cells with WT PilY1 could suggest a built-in property that allows for inherent heterogeneity in surface adaptation. That is, transient changes in PilY1 conformation (the spike signatures) may not be sufficient to drive signaling; only sustained conformational changes (i.e., plateaus) can do so. Our observation that the vWA-Cys152S mutant variant of PilY1 is skewed ~90:10 towards spike signatures (i.e., transient conformational changes), and that this strain is defective for c-di-GMP signalling, supports this conclusion. Thus, not every interaction between a cell and the surface to which it might attach is “productive”, a conclusion consistent with several reports showing the heterogenous nature of *P. aeruginosa* populations transitioning to a biofilm lifestyle [20, 38–40]. Furthermore, we could predict then that a PilY1 mutant that favors the plateau conformation should promote c-di-GMP signaling and be a hyper-biofilm former. We have performed extensive genetic screens looking for PilY1 mutants with such phenotypes with no success to-date. Thus, an alternative explanation is that the ability of TFP/PilY1 to transition *between* conformations is key to the ability to signal properly, and that locking the protein in one conformation, or another, results in aberrant signaling.

The critical role for vWA-Cys152 in c-di-GMP signaling and biofilm formation is supported by our genomic analysis, which highlight differences in the PilY1 protein among PA14 and PAO1 strains. The vWA domain of PilY1 from the PA14 and PAO1 strains are very different, with PilY1 proteins from the PAO1 clade (PAO1 and IPCD83) lacking the conserved vWA-Cys152 and vWA-Cys147 residues. *P. aeruginosa* PA14 strains engineered to carry the PAO1 or IPCD83 alleles of PilY1, which lack the conserved vWA-Cys152 and vWA-Cys147 residues, result in a hybrid strain that behaves very much like the *P. aeruginosa* PA14 strain carrying the vWA-Cys152S mutant protein. Thus, our genetic analysis confirmed that the observed sequence differences have functional consequences. The distinct PilY1 proteins of *P. aeruginosa* PA14 and PAO1 may also contribute to explaining the differences in the surface commitment strategies observed for these strains, as reported previously [3, 39].

Our AFM data show that force curve plateaus can be maintained for up to 50 ms; it is important to note that this value may be an underestimation because desorption of the pilus from the AFM tip may result in the force curve returning to baseline. With this important caveat in mind and considering that the *P. aeruginosa* TFP has a known retraction rate of ~0.5 μm s^−1^ [41], then the distance that the TFP is retracted during this 50 ms window (the time spane plateaus are maintained) is ~0.025 μm. This is quite a short distance (TFP can exceed two microns) and corresponds to the pilus being (almost) fully retracted, with the priming complex (i.e., the minor pilins) and the vWA domain of PilY1 docked into the pore of the secretin [5]. Furthermore, if we postulate that TFP/PilY1-mediated signaling is a consequence of mechanical force, for the TFP/PilY1 to remain under force and thus potentially capable of propagating a signaling event via a conformational change, we hypothesize that at least one other pilus would need to be bound to the surface to pull in opposition to the fully retracted pilus described above. That is, PilY1-mediated signaling would require multiple pili to decorate the cell surface – this model has a key corollary in that TFP must be robustly expressed for signaling to occur. Interestingly, previous studies [42–45] and work from our team [3] shows that the level of TFP is low in planktonic cells. Furthermore, our team showed that increased cAMP levels via the Pil-Chp system [3], which is key for pilus production, might require several cellular generations and multiple transient surface interactions to occur [39]. Thus, our previous observations of a role of multigeneration cAMP signaling via TFP may be *necessary* to produce multiple TFP; multiple TFP, in turn, are *required* for subsequent c-di-GMP signaling.

Based on the data presented here and previous studies from our team and others [3, 20, 39], we propose the following model of the early events initiating biofilm formation by *P. aeruginosa* PA14 (**Fig. 5**). When the TFP of *P. aeruginosa* PA14 initially engage the surface, we propose that the Pil-Chp signaling cascade promotes cAMP production, which in turn enhances transcription and subsequent production of TFP over the low levels of these appendages produced by planktonically-grown bacteria [3]. Currently, we do not have a strong mechanistic understanding of the linkage(s) among TFP, the Pil-Chp system and cAMP production. However, once more pili are deployed to the surface, this event provides the necessary condition for multiple surface-engaged TFP working in opposition to generate mechanical tension. This mechanical tension in turn can drive the sustained, PilY1-Cys152-dependent conformational changes we have observed for WT cells. That is, the conformational change in vWA domain of PilY1 is maintained only during the application of force when the TFP pull against a solid surface and thereby generate tension (with the cells presumably not moving). We propose that when multiple TFP are engaging the surface, the change in TFP/PilY1 conformation can be sustained as the pilus retracts and PilY1 is docked in the PilQ pore; here PilY1 can interact with PilO, as has been reported for the homologous system in *Myxococcus xanthus* [5]. Based on our recent study [20], the proposed PilY1-PilO interaction can in turn drive the documented PilO-SadC signal transduction cascade [20], which stimulates c-di-GMP signaling and increased biofilm formation. It is also important to note that a recent pull-down analysis indicate that there is one molecule of PilY1 per pilus in *M. xanthus* [5], thus it is unlikely that intermolecular disulfides are being formed with other PilY1 proteins. Additionally, cryo-electron tomography shows the C-terminal domain of PilY1 to be in direct contact with the minor pilins while the vWA domain is at the apex of the pilus fiber [5], suggesting that intermolecular disulfide bond formation between PilY1 and any of the minor pilins is also unlikely. Consistent with this conclusion, our purification of the vWA domain and Western analysis of cell-surface PilY1 shows no evidence of PilY1 forming intermolecular multimers.

**Figure 5.**
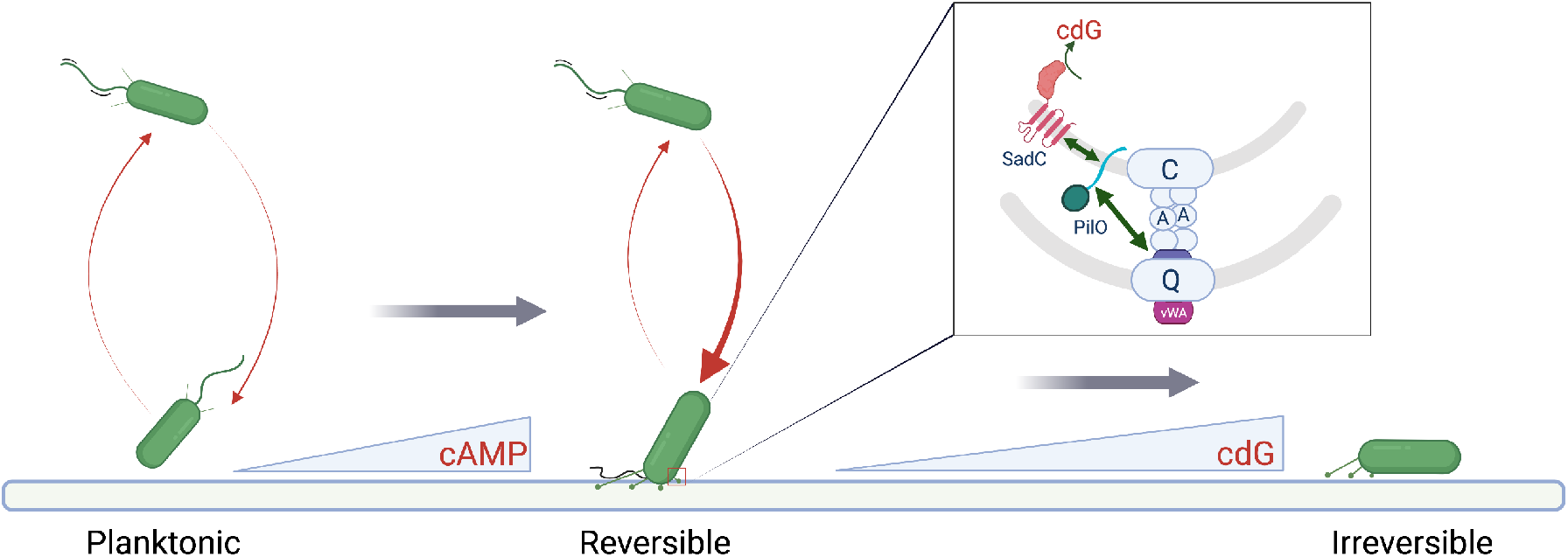
Proposed model for force-induced mechanical force drive transition from planktonic to irreversible attachment. Planktonic bacteria interact with the surface and increase cAMP levels and surface pili levels. The proposed PilY1-PilO interaction can in turn drive the documented PilO-SadC signal transduction cascade which stimulates c-di-GMP signaling and increased biofilm formation.

Finally, our studies were able to successfully separate the role of TFP in motility from its role in signaling. Work from our team and others [2, 3, 18, 20, 46] have implicated TFP in surface sensing via the surface-dependent stimulation of the second messengers cAMP and c-di-GMP, however, in these studies the mutants used also disrupted TFP-mediated twitching motility. Here, the Cys152S allele of PilY1 results in a clear loss of c-di-GMP signaling but strains carrying this mutation display robust twitching motility. Together with our previous studies showing a role of the TFP alignment complex component PilO in c-di-GMP production [20], we believe it is quite clear that TFP are not only a key appendage for adhesion and surface motility, but also a central player in surface-specific signal transduction.

## Materials and Methods

### Bacterial strains, plasmids, media and growth conditions

All bacterial strains used in this study are listed in the supplementary material in Table S1. *P. aeruginosa* PA14, *E. coli* S17-λ-pir were routinely grown in 5 mL lysogeny broth (LB) medium or struck on 1.5% LB agar plates with appropriate antibiotics, if necessary. Overnight cultures were grown in LB at 220 rpm on a roller drum. *Saccharomyces cerevisiae* InvSc1 (Thermo Fisher) used for cloning was maintained on yeast peptone dextrose (YPD - 1% Bacto yeast extract, 2% Bacto peptone, and 2% dextrose) with 2% agar. Synthetic defined medium without uracil (Sunrise Science Products) was used to select for yeast with construct. All chromosomal point mutation were constructed using the pMQ30 shuttle vector while pMQ72 multi-copy plasmid was used for ectopic expression. All plasmids and oligonucleotides used in this study are listed in Table S2 and Table S3 respectively.

### Construction of deletion mutant strains

All chromosomal in-frame gene deletions were constructed using the pMQ30 shuttle vector carrying the flanking regions of the gene via homologous recombination using the yeast machinery [47] or by Gibson cloning as previously described in [48]. For yeast cloning, *S. cerevisiae* was grown overnight at 30 °C in YPD. Synthetic defined medium without uracil (Sunrise Science Products) was used to select for yeast colonies with the plasmid construct. Plasmids were extracted from yeast using the ‘smash and grab’ method and transformed by electroporation into *E. coli* S17 cells and grown on LB plates with 10 μg/ml gentamycin at 30 °C overnight [2]. Colony polymerase chain reaction (PCR) amplification and sequencing was used to confirm plasmid construction. Plasmids were introduced in *P. aeruginosa* by conjugation and merodiploids were selected on 25 μg/ml gentamycin and 20 μg/ml nalidixic acid after which cells were counter-selected on LB with 10% sucrose-containing medium with no added salt [3]. Deletions were confirmed by colony PCR amplification and sequencing with primers flanking the gene. All sequencing was done at the Molecular Biology Core at the Geisel School of Medicine at Dartmouth.

### Construction of chromosomal point mutations

Point mutations in the *pilY1* gene were made using a modified *in vitro* site-directed mutagenesis protocol [49]. Forward and reverse complementary primers consisting of the nucleotide codon sequence encoding for the mutation of interest were used to separately amplify the pMQ30 (for chromosomal mutations) or pMQ72 (ectopic expression) parental plasmids with the gene of interest using high fidelity Phusion polymerase (NEB). After four cycles of amplification, the products of these reactions were combined and amplified for an additional 18 cycles with additional Phusion polymerase added. The parental plasmid was digested for 4 h using Dpn1 endonuclease (NEB) at 37 °C. Following digestion, the PCR product was transformed into *E. coli S17* competent cells and selected on LB with 10 μg/ml gentamycin. Plasmid containing the desired point mutation was isolated and confirmed by sequencing. Introduction of mutations on the chromosome was done by conjugation and counter-selection as described above. All chromosomal mutations were verified by PCR amplification and sequencing.

### Biofilm assay

Overnight cultures (1.5 μl) were inoculated in U-bottom 96 well plates (Costar) containing 100 μl M8 salts minimal medium (Na_2_HPO_4_, KH_2_PO_4_ NaCl) supplemented with glucose (0.2% v/v), casamino acids (0.5% v/v) and MgSO_4_ (1 mM), subsequently referred to as M8 medium. Biofilm assay plates were then stained with 100 μl of 0.1% crystal violet in water for 20 mins at room temperature and destained for 20 mins with 125 μl de-staining solution (40% glacial acetic, 40% methanol and 20% H_2_O v/v). Absorbance was read at OD550 and destaining solution was included as the blank. Biofilm assays were done similar to published work by the O’Toole group [50, 51].

### In vivo c-di-GMP quantification

Nucleotides were extracted from *P. aeruginosa* cells scraped from 0.52% agar with M8 medium after incubation for 37°C for 14 h. Cells were removed from plates by gently scraping with a cover slip to avoid scraping the agar, and then immediately placed on ice. Cell pellets were re-suspended in 250 μl nucleotide extraction buffer (methanol/acetonitrile/dH_2_O 40:40:20 + 0.1 N formic acid) and incubated at −20°C for 1 h. Following nucleotide extraction, cells were spun for 5 mins at 4°C, 200 μl of supernatant was removed and then added to 8 μl of 15% NH_4_HCO_3_ stop solution. Nucleotides were dried in a speed vacuum and resuspended in 200 μl HPLC grade water (JT Baker) and placed in screw cap vials (Agilent Technologies). Quantification of c-di-GMP levels was done by liquid chromatography-mass spectrometry (LC-MS/MS) by Lijun Chen at the Mass Spectrometry Facility at Michigan State University. All samples were normalized to dry weight and expressed as 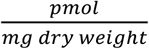.

### Macroscopic twitch assay

One percent LB agar plates were stab inoculated using toothpicks dipped in overnight cultures to the bottom of the plate. Plates were incubated at 37°C for 24 h and an additional day at room temperature. The agar was subsequently removed from the petri plate and the twitch zones stained with crystal violet to visualize, and the diameter of the twitch zones measured.

### Plaquing assay

One percent agar plates (60 x15 mm) with M8 medium were prepared and cooled to room temperature. Fifty microliters of *P. aeruginosa* overnight culture were added to 1 mL of 0.5% warm top agar made with M8 medium and gently mixed. The mixture was quickly poured onto 1% agar plates made with M8 salts. Plates were swirled to ensure even spreading of top agar. Once cooled, 2 μl of phage DMS3_vir_ strain were spotted to the center of the plate, allowed to dry and subsequently incubated at 37 °C overnight.

### Cell surface pili

WT, Δ*pilY1, vWA* variants and vWA-Cys152S cells were streaked in a grid-like pattern on 10% agar plates with M8 SALTS/supplements and incubated at 37 °C overnight. Four plates per strain were struck for each biological replicate to ensure adequate number of pili could be recovered. The following day cells were scraped off the plate using a cover slip and put in a 2 mL tube and vortexed vigorously for 2 mins with 1 mL of phosphate buffer saline (PBS – Corning). Cells suspensions were subsequently spun at 16, 000 x g for 5 mins in a table-top centrifuge and the supernatant removed and transferred to a clean tube and spun again. This step was repeated until no pelleted cells were recovered. Proteins were precipitated with 20% trichloroacetic acid (TCA – VWR) on ice overnight at 4° C. Precipitated proteins were collected by centrifuging at 16 000 x g for 25 mins at 4° C. The supernatant was discarded and the tubes re-centrifuged for 3 mins to get rid of any remaining supernatant. Pellets were washed twice with 1 mL acetone (VWR) and subsequently air dried to remove residual acetone. Pellets were resuspended in 100 ml 1x sample buffer (BioRad) with b-mercaptoethanol and boiled for 5 mins and then briefly spun before being ran on a 12% SDS-PAGE gel, and the samples probed for PilA and FliC by Western blot analysis. FliC served as the loading control and was used for normalization of PilA protein levels. Samples were also resolved on a 7.5% SDS-PAGE gel and probed for PilY1 using a-PilY1 antibody generously provided by Matt Wolfgang. Western blot analysis was performed as described below.

### Western Blot analysis for PilY1 protein levels in whole cell lysate

All strains were grown overnight in LB at 37 °C. For whole cell lysate (WCL) preps, overnight cultures were diluted 1:100 in 5 mL M8 SALTS/supplements minimal medium and sub-cultured for ~3 h at 37 °C. Samples were resolved on a 7.5% Tris-HCl precast SDS-PAGE gel (Bio-Rad) and blotted onto 0.45 μm pore size nitrocellulose membrane (Bio-Rad) using the 1.5 mm pre-programmed method on a Trans-Blot Turbo Transfer System (Bio-Rad). The membrane was incubated in blocking buffer (LI-COR Blocking Buffer in TBS) for 1 h at room temperature and incubated for 1 h or overnight at 4 °C in polyclonal a-PilY1 antibody (1:5000 dilution) in BSA TBST_0.1%_ buffer. Following incubation with primary antibody, the membrane was washed in TBST_0.1%_ for 5 mins x3 and incubated for 1 h with goat anti-rabbit in TBST_0.1%_ (1:10,000 dilution) secondary antibody (LI-COR IRDye® 800CW Goat a-rabbit). Incubation with secondary antibody and all subsequent steps were performed in the dark. After incubation with the secondary antibody, the membrane was washed in TBST_0.1%_ for 5 mins x2 and then once in TBS. The membrane was imaged using the LI-COR Odyssey CLx imager at BioMT Core at the Geisel School of Medicine at Dartmouth. PilY1 protein levels were quantified relative to the cross-reacting band at ~60 kD using the LI-COR Image Studio Lite software by drawing a rectangle of the same size around each band and using the following background settings: average, border width of 3, segment = all.

### Protein quantification

Total protein levels in whole cell lysate was quantified using the Bio-Rad protein assay Dye Reagent as per the manufacturer’s instructions as outlined by Bradford [52].

### AFM force spectroscopy (AFM)

Overnight cultures used in AFM experiments were diluted 200-fold in M8 salts and seeded on hydrophobic non-treated polystyrene petri dishes (Corning) and left for 10 minutes to adhere [25]. Dishes were then washed gently but thoroughly with M8 salts medium to remove most non-adhered bacteria and used for AFM experiments in the same medium. AFM experiments were performed at room temperature using a NanoWizard® 4 NanoScience AFM (JPK Instruments). Gold cantilevers (PNP-TR probes – Pyrex Nitride Probe with Triangular Cantilevers – from NanoWorld) were treated for 16 h with a 1 mM 1-dodecanethiol solution in ethanol to render them hydrophobic, then rinsed with ethanol and kept in milliQ water until AFM experiments were ready to be performed. Prior any measurements, the cantilever’s spring constant was empirically determined by the thermal noise method [53]. The AFM force volume mode was used to record force-distance curve in a pixel-by-pixel manner (force mapping) on 6 × 6 μm^2^ areas (32 × 32 pixels, i.e. 1024 curves) with a bacterium at the center, previously localized by an optical microscope coupled to the AFM. For the Δ*pilA* strains overexpressing WT PilY1 or PilY1-Cys152S and lacking the pilus fiber, the area was decreased to 1 μm_2_ around the bacterium’s poles. The following settings were used: an applied force of 250 pN, a constant approach/retract speed of 5 μm/s and a z-range of 1.5 μm.

### AFM data analysis

Data were analyzed with the data processing software from JPK Instruments (Berlin, Germany). In a first approach, all force distance curves exhibiting an adhesive event were selected, as opposed to the non-adhesive curves which were discarded, thus allowing an estimation of the binding probability. In a second approach, adhesive curves were sorted depending on their signature (plateaus vs spikes) and the maximum adhesion sustained by each adhesive peak was determined. The frequency of plateaus was assessed by dividing the number of curves showing plateaus plus curves showing both plateaus and spikes by the total number of adhesive curves. A similar approach was used to calculate the percent of spikes. The formulas to calculate the percent plateaus (P_plateaus_) or the percent spikes (P_spikes_) are shown below:

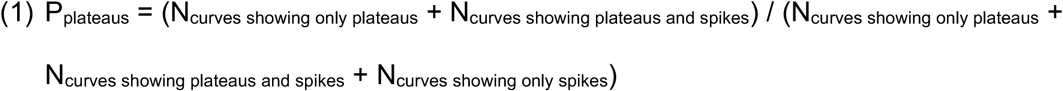

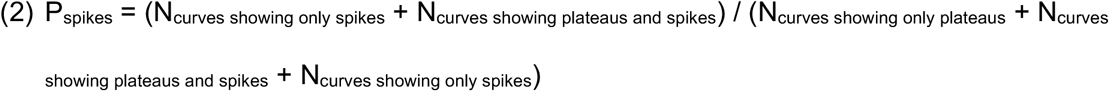

Statistical analyses were performed with Origin.

### Analysis of IPCD database: generation of phylogenetic tree, alignment and calculation of Shannon Diversity

We performed nucleotide BLAST searches on a local version of the IPCD database of *P. aeruginosa* genomes to identify variants of the PilY1 protein. Using the nucleotide sequences of PilY1 from PA14, PAO1 and IPCD-83 (GenBank: MCMY00000000), we were able to identify 852 strains with versions of the full protein. We used custom MATLAB scripts to perform an alignment of the amino acid sequences of all 852 versions of PilY1 and construct the corresponding phylogenetic tree. We performed the alignment of PilY1 amino acid sequences using a series of BLOSUM80 to BLOSUM30 scoring matrices. We constructed the phylogenetic tree of PilY1 sequences using a Jukes-Cantor maximum likelihood method to estimate the number of substitutions between two sequences and an Unweighted Pair Group Method Average (UPGMA) to construct the phylogenetic tree from the pairwise distances. 129 sequences of PilY1 belong to a clade with highly similar proteins, which includes PilY1 from PA14. 723 sequences belong to a diverse clade that includes PilY1 from PAO1 and IPCD-83. Within each of these two groups, we calculated the Shannon diversity in each position along the aligned amino acid sequence using *H* = −∑ *pi* ln (*pi*), where *pi* is the probability of each amino acid (including gaps). Code is available at github.com/GeiselBiofilm.

### Growth assays

Overnight cultures were inoculated in M8 salts/supplements at a starting OD600 of ~0.05. Readings were taken every 40 mins for 16 h in a Synergy Neo2-multimode microplate reader at the BioMT Core at the Geisel School of Medicine at Dartmouth.

### Cloning and protein expression of GST-vWA fusions

The coding region of the WT and the C152S mutant of the vWA domain (amino acids 30-369) from *P. aeruginosa* PA14-UCBPP were PCR amplified and cloned into pGEX-6p-1 plasmid at the BamHI cut site by Gibson assembly. *E. coli* BL21 (DE3) competent cells were transformed with plasmid and selected on LB plates with 50 μg/mL carbenicillin grown at 30 °C overnight. A single colony was used to inoculate 5 mL of liquid LB and grown for 12-14 h at 30 °C. Each 5 mL seed culture was used to inoculate 500 mL LB in a 2 L flask and allowed to grow at 37 °C with shaking 200 rpm until the OD600 was 0.6-0.8. A total of 6 L LB (12 flasks) were inoculated. Expression was induced with 0.1mM isopropylβ-D-1-thiogalactopyranoside (IPTG) for 4 h at 37 °C. Bacteria were harvested at 5,000 × *g* for 10 min, washed with PBS buffer and stored at −20 °C until further use.

### Purification of wild-type GST-vWA and Cys152S mutant proteins

*E. coli* cells expressing WT GST-vWA and GST-vWA-C152S mutant proteins were resuspended in PBS supplemented with 2 mM TCEP (Thermo scientific), 0.01 mg/mL lysozyme from chicken egg (SIGMA), EDTA-free protease inhibitors cocktail (BImake) and 10U/mL benzonase nuclease (Millipore) and lysed in a Microfluidizer LM10 (Microfluidics) at 18,000 psi. Nucleic acids were precipitated by addition of 0.1% polyethylenimine branched (SIGMA). Crude cell lysates were cleared by centrifugation at 200,000 × *g* for 1 hour at 4 °C in a Beckman Optima L-70 ultracentrifuge. Clear lysates were incubated overnight with 5 mL Glutathione Sepharose 4B resin (Cytiva) previously equilibrated with PBS containing 2 mM TCEP. Lysates and resin were transferred to a disposable plastic column and allowed to drain fully (flow through). Resin was washed with at least 15 column volumes of PBS, 2 mM TCEP buffer before elution of the GST-vWA proteins with 5 column volumes of freshly prepared elution buffer 50 mM Tris -HCl pH 8, 10 mM reduced glutathione. Elution fractions were concentrated using 30,000 MWCO 15 mL Amicon centrifugal filters (Millipore) in a Beckman Allegra 6R centrifuge. Proteins were loaded in a HiLoad Superdex 75 pg (Cytiva) pre-packed column equilibrated with 50 mM Tris-HCl pH 8, 150 mM NaCl, 1 mM TCEP using an AKTApure instrument (Cytiva). Fractions containing the fusion GST-vWA protein were combined and concentrated as before and subjected to a second gel filtration step using a high-resolution Superdex 200 increase 10/300 (Cytiva). Purified WT and C152S mutant GST-vWA proteins were extensively dialyzed against 20 mM sodium phosphate pH 7.4 buffer Final protein concentrations were determined using Bio-Rad protein assay reagent.

### Circular Dichroism (CD) and melting curves

The far-UV circular dichroism (CD) spectra (195–250 nm) were recorded with a JASCO J-815 spectrophotometer (Jasco, Inc.) equipped with a CDF426S/15 Peltier temperature controller using a 2-mm path length quartz cuvette. CD spectra of proteins were recorded at 20 °C using a step size of 0.1 mm. A time constant of 12 s was used to improve the signal to noise ratio and to decrease the contribution of the solvent at lower wavelengths. CD spectra were recorded using 1 μM of GST-vWA WT and GST-vWA-C152S proteins in 20 mM sodium phosphate buffer, pH 7.4, and corrected by subtracting the spectrum of the buffer alone.

Thermal unfolding curves were obtained by monitoring the ellipticity at 222 and 208 nm of both fusion proteins at 1 μM concentration at a heating rate of 1 °C min^−1^ in the temperature range of 20 to 90 °C. A 1s integration time and 5s equilibration time were used for each measurement and buffer ellipticities at the selected wavelengths were subtracted from the samples data. Raw CD data were converted into the molar ellipticity [θ]_λ_ (deg cm^2^ dmol^−1^) at each wavelength using the relation, [θ]_λ_ = θ_λ_/(10CNl), where θ_λ_ is the observed ellipticity in millidegrees at wavelength λ, *C* is the molar protein concentration, *N* is the number of amino acids of the protein, and l is the path length of the cuvette in cm. Following CD measurements, protein samples were collected, and protein concentrations measured for accuracy.

## Acknowledgments

We thank Roger Levesque for providing the IPCD strains used in this report and Dr. Sherry Kuchma for building the Δ*vWA_p_* strain. Also, thanks to Dr. Matt Wolfgang for the PilY1 antibody, Emilie Shipman for help with protein purification, Dr. Paul Delfino for technical assistance with CD, Kelsie Leary, Dr. Dean Madden and Dr. Holger Sondermann for advice on analyzing the CD data. The authors would also like to thank rotation students who worked on the project: Alexander Pastora and Rebecca Valls, other members of the O’Toole lab: Chris Geiger, Dr. Sherry Kuchma and Fabrice Jean-Pierre for helpful discussions. This work was supported by the NIH via awards R37 AI83256 (to G.A.O.), R01 AI43730 (to G.C.L.W.) and COBRE/NIGMS 5 P20 GM130454-02 (to D.S). This work was also supported by bioMT through NIH NIGMS grant P20-GM113132. Work at UCLouvain was supported by the Excellence of Science-EOS programme (Grant #30550343), the European Research Council (ERC) under the European Union’s Horizon 2020 research and innovation programme (grant agreement n°693630), and the National Fund for Scientific Research (FNRS). Y.D. is a Research Director at the FNRS.

